# Partitioning gene-mediated disease heritability without eQTLs

**DOI:** 10.1101/2021.07.14.452393

**Authors:** Daniel J. Weiner, Steven Gazal, Elise B. Robinson, Luke J. O’Connor

**Affiliations:** Stanley Center for Psychiatric Research, Broad Institute of MIT and Harvard, Cambridge, MA, USA; Program in Medical and Population Genetics, Broad Institute of MIT and Harvard, Cambridge, MA, USA; Harvard T.H. Chan School of Public Health, Harvard University, Boston, MA, USA; Keck School of Medicine, University of Southern California, Los Angeles, CA, USA

## Abstract

Unknown SNP-to-gene regulatory architecture complicates efforts to link noncoding GWAS associations with genes implicated by sequencing or functional studies. eQTLs are used to link SNPs to genes, but expression in bulk tissue explains a small fraction of disease heritability. A simple but successful approach has been to link SNPs with nearby genes, but the fraction of heritability mediated by these genes is unclear, and gene-proximal (vs. gene-mediated) heritability enrichments are attenuated accordingly. We propose the Abstract Mediation Model (AMM) to estimate (1) the fraction of heritability mediated by the closest or *k*^*th*^-closest gene to each SNP and (2) the mediated heritability enrichment of a gene set (e.g. genes with rare-variant associations). AMM jointly estimates these quantities by matching the decay in SNP enrichment with distance from genes in the gene set. Across 47 complex traits and diseases, we estimate that the closest gene to each SNP mediates 27% (SE: 6%) of heritability, and that a substantial fraction is mediated by genes outside the ten closest. Mendelian disease genes are strongly enriched for common-variant heritability; for example, just 21 dyslipidemia genes mediate 25% of LDL heritability (211x enrichment, P = 0.01). Among brain-related traits, genes involved in neurodevelopmental disorders are only about 4x enriched, but gene expression patterns are highly informative, with detectable differences in per-gene heritability even among weakly brain-expressed genes.

## Introduction

A common challenge in GWAS is to map disease-associated variants to the target genes that mediate their effects. Associated loci often contain multiple genes^1,2^, fine-mapped causal variants are mostly noncoding,^3,4^ and functional data, including from eQTLs, is often inconclusive.^5–7^

Conversely, it is often unknown which SNPs regulate putative disease genes. Disease genes with large-effect rare variants sometimes localize near common-variant associations from GWAS^2,8,9^, suggesting possible mediation, but unknown SNP-to-gene architecture complicates mediation analyses. Window-based enrichment approaches (like LDSC-SEG and MAGMA) can be used to show that a set of genes is enriched for nearby SNP associations.^10–15^ However, the magnitude of these window-based enrichments is attenuated compared with the mediated heritability enrichment of the genes themselves.

A different approach is to use eQTL data, since eQTLs can identify the set of SNPs regulating expression of a nearby gene.^16–21^ However, eQTL data may fail to recapitulate disease-relevant cell types and cellular states, which are often unknown. While measured expression from bulk tissue is more readily available, a recent study estimated that the fraction of heritability mediated in *cis* by tissue-level gene expression is only 11%.^22^ Moreover, colocalization analyses suggest that eQTLs and nearby GWAS loci usually arise from different causal SNPs.^23^

We propose the *Abstract Mediation Model* (AMM) for the relationship between SNPs, genes and a disease or complex trait. Under the model, the effect of a disease-associated SNP is mediated by one or more nearby genes:

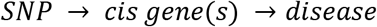

where the genes in this model are abstractions for whatever biological variables actually have a causal effect on disease risk (for example, expression levels in a certain cell type). Unlike in eQTL studies, these variables are not observed, and neither are the SNP-to-gene or gene-to-disease effect sizes. Instead, we observe proxies for both effects: the proximity of SNPs to genes, and an annotation for each gene, such as its membership in a disease-relevant gene set (**Figure 1A**). This limited information is sufficient to partition mediated heritability across genes. In particular, it allows us to estimate (1) the fraction of heritability mediated by the *k*^*th*^ closest gene to each SNP, and (2) the proportion of heritability mediated by a specified gene set.

**Figure 1.**
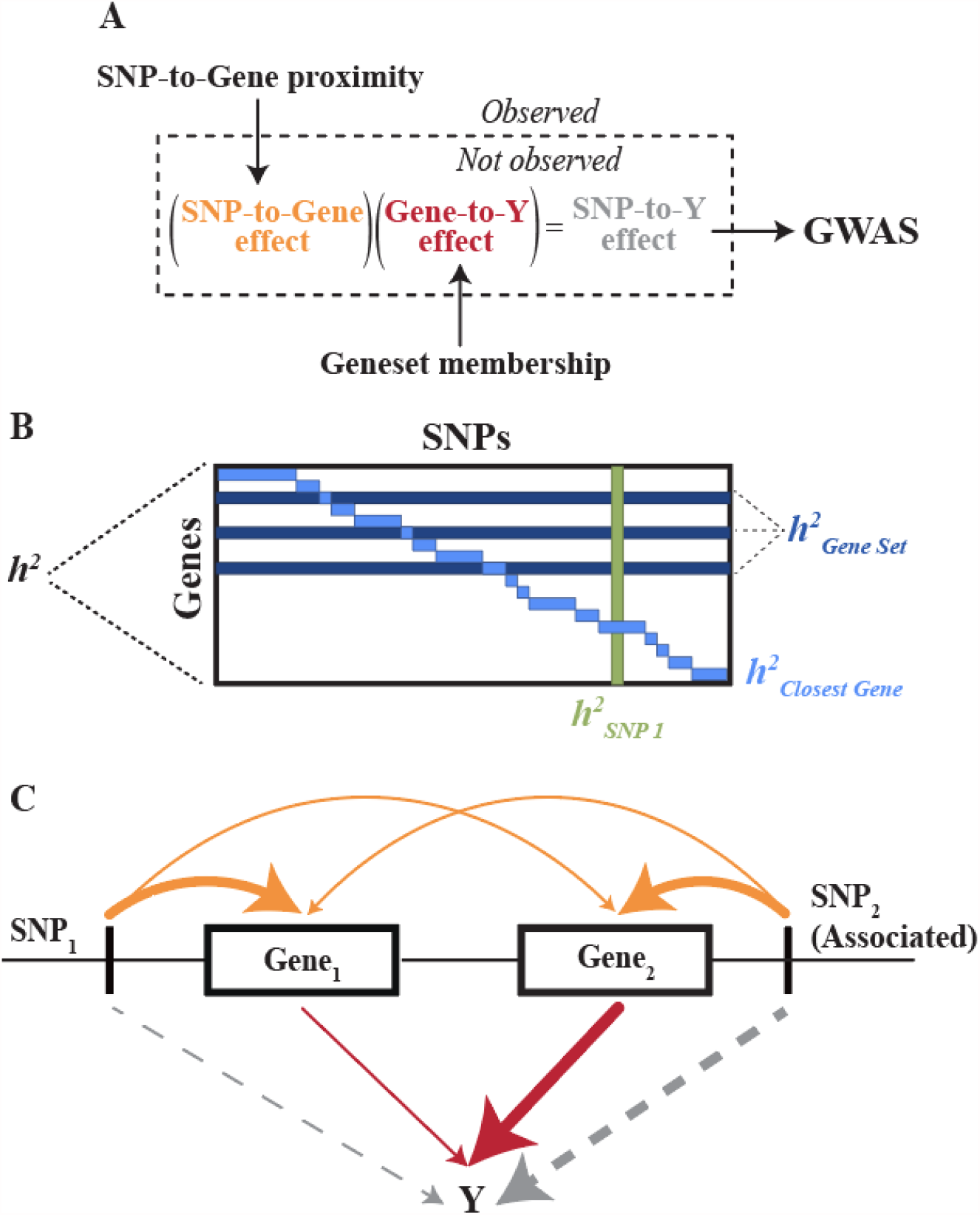
The Abstract Mediation Model. **(A)** Under AMM, the gene-mediated effect of a SNP on a trait is equal to the product of the SNP-to-gene and gene-to-trait effects. Neither of these quantities is observed directly, but we do observe proxies for each: SNP-to-gene proximity, and geneset membership, respectively. **(B)** AMM partitions mediated heritability across SNP-gene pairs, each an entry in the SNPs-by-genes matrix. The heritability mediated by a gene set is a row sum (dark blue horizontal rectangles), and similarly, the heritability mediated by the closest gene to each SNP is the sum across SNP-closest-gene pairs (blue diagonal cells). **(C)** Example of mediated heritability enrichment under AMM. The orange arrows represent SNP-to-gene effect sizes, *p*^*(k)*^; larger values of *p*^*(k)*^ are denoted with arrow thickness. The red arrows represent gene-to-trait effect sizes. The gray arrows represent SNP-to-trait effect sizes. SNP_2_ has a large effect on trait Y (thick gray arrow) because its closest gene (Gene_2_) is a member of an enriched gene set and has a large effect on Y (thick red arrow).

## Results

### Intuition

On average across the genome, some fraction of heritability is mediated by the closest gene to each SNP, denoted *p*^*(1)*^. Suppose that every disease-associated SNP regulated its closest gene, such that 100% of heritability was mediated by the closest gene to each SNP (*p*^*(1)*^ = 1). Under this model, the proportion of heritability mediated by a set of genes would be equal to the heritability explained by the SNPs whose closest gene is in that gene set.

More generally, suppose that we knew the proportion of heritability mediated by the closest, second-closest, and *k*^*th*^-closest gene. Even without knowing the specific disease gene(s) for individual disease-associated SNPs, we could estimate the proportion of heritability mediated by a set of genes: the heritability of SNPs whose *k*^*th*^ closest gene is in the gene set, multiplied by *p*^*(k)*^, summed across k. That is, if the SNP-to-gene architecture were known, it would allow us to estimate the proportion of heritability mediated by a set of genes.

Conversely, suppose that *p*^*(k)*^ were unknown, but that we did have a set of genes known to be enriched for heritability (like constrained genes; see below). The set of SNPs whose *k*^th^-closest gene is in the gene set would be enriched for heritability, and this enrichment would be proportional to *p*^*(k)*^. For example, if SNPs are three times as likely to affect their closest gene compared with their second, then SNPs adjacent to a known disease gene will have triple the heritability enrichment as those one gene away. By comparing the enrichments of gene-set-proximal SNPs with that of the gene set itself, we would obtain estimates of ^(*k*)^.

### The Abstract Mediation Model

The Abstract Mediation Model (AMM) partitions heritability across SNP-gene pairs. The genes in the model are abstractions for whatever genic variables actually modulate disease risk (mRNA levels, enzyme activity, etc.), hence “abstract”; the heritability assigned to a set of SNP-gene pairs is referred to as *mediated heritability*. The expected mediated heritability of a SNP-gene pair is equal to the SNP-to-gene effect-size variance times the gene-to-trait effect-size variance (**Figure 1A**). The SNP-to-gene effect-size variance depends on whether that gene is the closest, second closest, or *k*^th^ closest gene to that SNP; the gene-to-trait effect-size variance depends on whether the gene is in an enriched gene set. The heritability mediated by a gene set is the sum of the mediated heritability across all SNP-gene pairs where the gene is in that set, and similarly, the heritability mediated by the closest gene to each SNP is the sum across SNP-closest-gene pairs (**Figure 1B**).

In detail, AMM jointly estimates the SNP-to-gene architecture (*p*^*(k)*^) and the mediated heritability enrichment of a gene set. We first train AMM *p*^*(k)*^ estimates on genes intolerant of heterozygous disruption (constrained genes), a large gene set enriched for common variant heritability across a range of traits.^24^ Specifically, the fraction of heritability mediated by the *k*^*th*^ closest gene, *p*^*(k)*^, is equal to the heritability of SNPs whose *k*^*th*^ closest gene is constrained, divided by the heritability of all SNPs with a *cis* constrained gene, conditioned on SNP-level functional annotations in the Baseline LD model (Online Methods).^25^ This estimated SNP-to-gene architecture is used to estimate the heritability enrichment of other gene sets, some of which are much smaller. The estimates of heritability enrichment of other gene sets are robust to the use of constrained genes to estimate *p*^*(k)*^ (Supplementary Figure 1). This approach allows for estimation of SNP-to-gene architecture without incorporating eQTL data.

If the gene set is enriched for heritability, then SNPs whose closest genes are in the gene set are more likely to be associated with the trait (**Figure 1C**). For a SNP *x*_*i*_ whose *k*^*th*^ ranked gene is in an enriched gene set A, its expected heritability is:

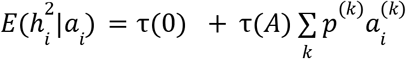

where τ(0) is the expected heritability of a SNP with no *cis* genes in annotation *A*, τ(*A*)is the excess per-SNP mediated heritability of genes in *A, p*^(*k*)^is the proportion of *cis*-mediated heritability explained by the *k*^th^-closest gene for each SNP, and 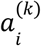 ∈ {0, 1}indicates whether the *k*^th^-closest gene to SNP *x*_*i*_ is in *A*. τ(*A*)can be interpreted as the additional effect-size variance for genes in *A* compared with the genome-wide average.

This expression leads to a stratified LD score regression equation^25^ (see Online Methods):

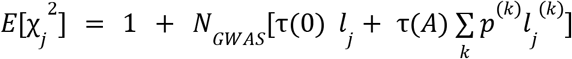

where 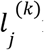 is the stratified LD score for SNP_j_ to the annotation a^(k)^, and *l* is the unstratified LD score of SNP *x*_*j*_.

The above regression can be used to estimate the heritability enrichment of gene set *A* (see Online Methods):

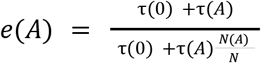

where *N(A)/N* is the fraction of genes in gene set *A*. If the gene set *A* is unenriched, then τ(*A*) = 0, and *e*(*A*) = 1. τ(*A*) can also be negative, in which case *A* is depleted. Finally, we define the fraction of heritability mediated by a gene set as the heritability enrichment times the fraction of genes in *A*:

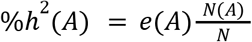

### Performance of AMM in simulations

We evaluated AMM in simulations with no LD. Stratified LD score regression, which we use to estimate fractions of heritability, is known to account for LD appropriately.^25,26^ We chose simulation parameters to approximately match UK Biobank (GWAS N=500,000; M=900,000 SNPs) (see Online Methods). We simulated 18,000 genes, 10% of which were assigned at random to an enriched gene set. SNP-to-gene and gene-to-trait effect sizes were drawn from point-normal distributions, with different parameters depending on SNP-to-gene proximity and on gene-set membership, respectively.

AMM produced unbiased estimates of *p*^*(k)*^, the fraction of heritability mediated by the *k*^*th*^ closest gene (**Figure 2A**). It also produced unbiased estimates of gene-set enrichment (**Figure 2B**). These simulations indicate that AMM can partition gene-mediated heritability without observing the genes themselves.

**Figure 2.**
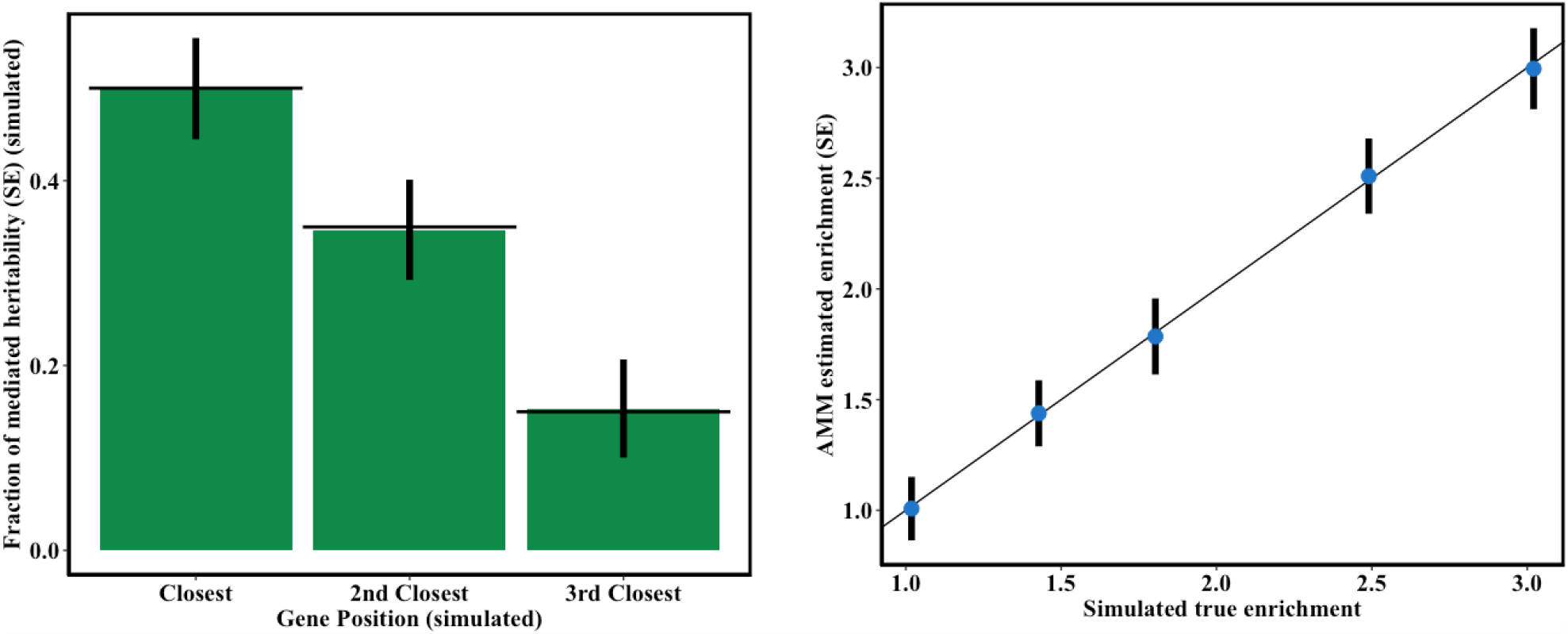
Performance of AMM in simulations. **(A)** AMM produces unbiased estimates of the proportion of heritability mediated by the *k*^*th*^ closest gene. Horizontal black segments are the mean of the true heritability proportion across simulation iterations. **(B)** AMM produces unbiased estimates of mediated heritability enrichment of gene sets. Error bars represent standard errors across 100 replicates.

### SNP-to-gene architecture of 47 complex traits

Constrained genes (pLI ≥ 0.9, n = 2,776, Supplementary Table 1), which are intolerant of heterozygous loss of function variation, are enriched for heritability across a wide range of complex traits.^24^ We applied AMM to this gene set in conjunction with well-powered GWAS summary statistics for 47 traits and common diseases (median N = 419,236) (Supplementary Table 2). We estimated the proportion of heritability explained by the closest, second-closest, and *k*^th^-closest gene to each SNP, and we meta-analyzed across traits (Online Methods).

On average, the closest gene to each SNP mediates 27.1% (SE: 6.4%) of heritability **(Figure 3A)**. This estimate is approximately concordant with eQTL data^20^ (Supplementary Figure 2) and epigenomic data.^17,27,28^ Per-gene heritability decays quickly from the closest gene to the third-closest, but detectable mediation persists as far as the 11-20th closest, where on average, each of the 11th through 20th closest genes mediate 2.3% (SE: 0.7%) of heritability.

**Figure 3.**
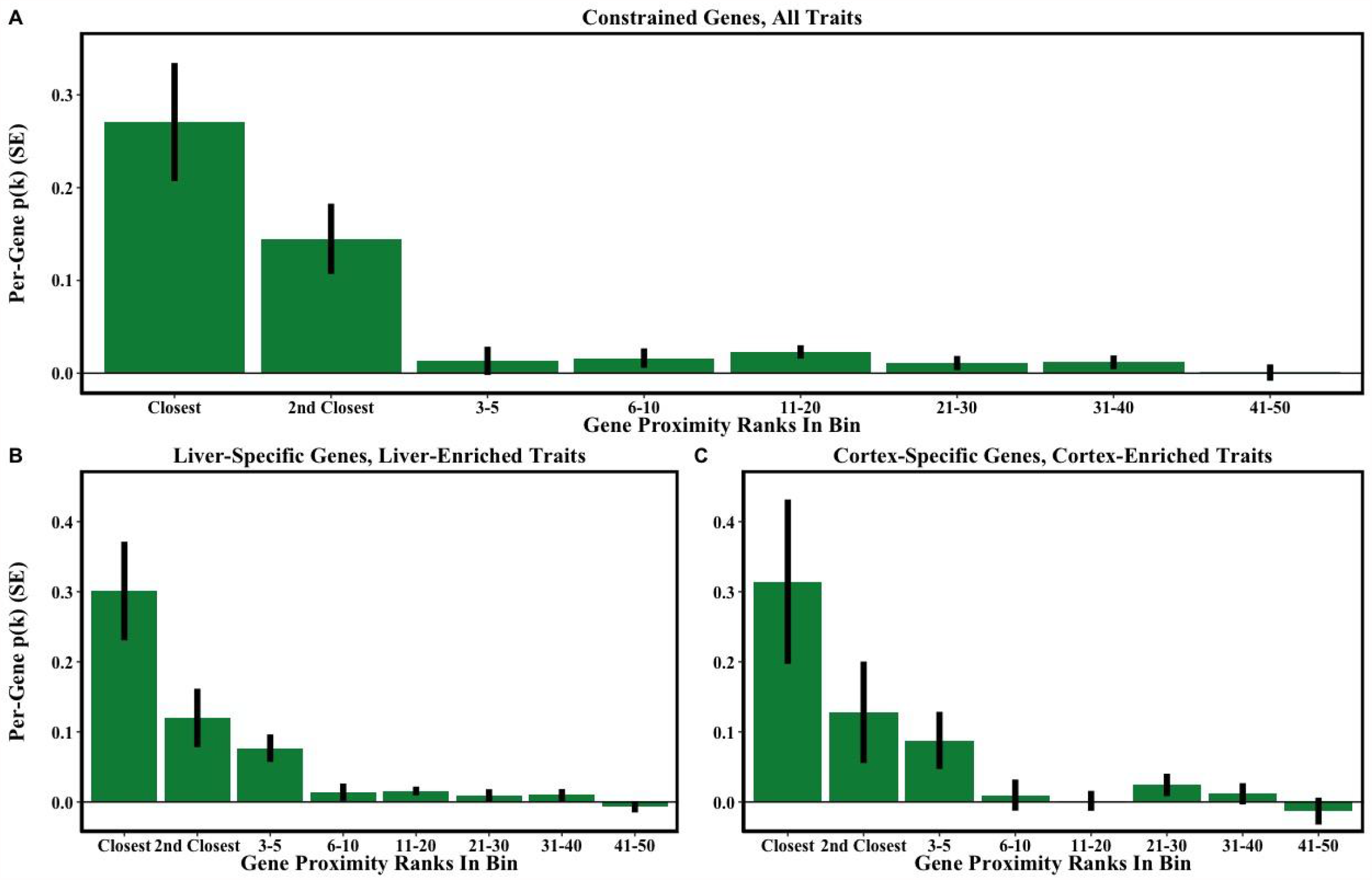
SNP-to-Gene architecture of 47 complex traits. **(A)** Mediated heritability across gene-proximity bins, meta-analyzed across 47 traits, estimated with the constrained gene set (pLI ≥ 0.9). The estimate of *p*^*(k)*^ is the average for genes in that bin; per bin *p*^*(k)*^ multiplied by the number of genes in the bin, summed across bins, equals 100% of heritability. Error bars represent standard errors. **(B)** As in (A), but using the set of genes specifically expressed in liver from GTEx (n = 1,766) and meta-analyzed across 8 traits with positive mediated enrichments in liver (Supplementary Table 4). (**C)** As in (A), but using the set of genes specifically expressed in cortex from GTEx (n = 1,766) and meta-analyzed across 9 traits with positive mediated enrichments in cortex (Supplementary Table 3). For numerical results, see Supplementary Table 5.

We replicated these results using previously-defined gene sets derived from tissue-specific gene expression data in GTEx.^10,20^ To maximize power in the meta-analysis, we chose two tissues, cortex and liver, that were enriched across several tissue-relevant traits (see below, and Supplementary Tables 3-4). Estimates of SNP-to-gene architecture were concordant: using the cortex gene set we estimate the closest gene mediates 31.4% (SE: 11.7%) of heritability, and using the liver gene set we estimate 30.1% (SE: 7.0%) (**Figure 3B-C**).

We performed three secondary analyses. First, we verified that the constrained gene set is broadly enriched across traits (median = 2.1x), confirming that it is appropriate to view our estimates as an average across traits (not only across a subset of enriched traits) (Supplementary Figure 3). Second, we considered the possibility that apparent signal in quite faraway genes could be driven by physical clustering of constrained genes, which could violate the no-spurious-enrichment assumption (see Online Methods); however, constrained genes do not cluster next to each other (and moreover, we jointly model all nearby genes in the regression) (Supplementary Figure 4). Third, we estimated *p*^*(k)*^ as a function of SNP-to-gene distance. We did not detect a statistically significant difference in SNP-to-gene architecture after incorporating distance (Supplementary Figure 5).

### Mediated heritability enrichment in Mendelian and drug target gene sets

Although most complex-trait heritability is explained by common regulatory variation, rare coding variants can have much larger effect sizes on these traits or on closely-related Mendelian phenotypes. It is unclear to what extent common-variant heritability is mediated by Mendelian disease genes.

We applied AMM to estimate common-variant enrichment in Mendelian gene sets associated with rare forms of common diseases or traits. We analyzed 21 Mendelian dyslipidemia genes (also including lipid drug targets)^29^, 50 Mendelian diabetes genes (also including drug targets)^29^, 206 skeletal growth disorder genes^2^, and 251 developmental disorder genes^30^ (predominantly neurodevelopmental) (Supplementary Table 1). None of these gene sets were defined using GWAS results (which would lead to bias; see Online Methods).

Mendelian gene sets were highly enriched for common-variant heritability (**Figure 4**). The 21 dyslipidemia genes mediate 25.1% (SE: 11.4%, P = 0.01) of common-variant heritability for low density lipoprotein (LDL), an enrichment of 211.5x (SE: 96.0x). By comparison, the set of constrained genes is 132 times larger, and it mediates approximately the same proportion of LDL heritability. For diabetes, a similar fraction of heritability is mediated by 50 Mendelian diabetes genes, representing a 64.4x enrichment (SE: 27.0x, P = 9.5e-3). These gene sets were not strongly enriched for height, but height enrichment was observed for skeletal growth disorder genes: we estimate that this gene set mediates 12.7% of common-variant heritability, a 10.9x enrichment (SE: 2.7x, P = 1.1e-4). There was a surprising 8.6x enrichment (SE: 2.7x, P = 2.3e-3) of neuroticism in the diabetes gene set; other brain-related traits were not enriched in this gene set (Supplementary Table 6), and S-LDSC is known to occasionally produce false positives for small annotations (see Discussion).^31^

**Figure 4.**
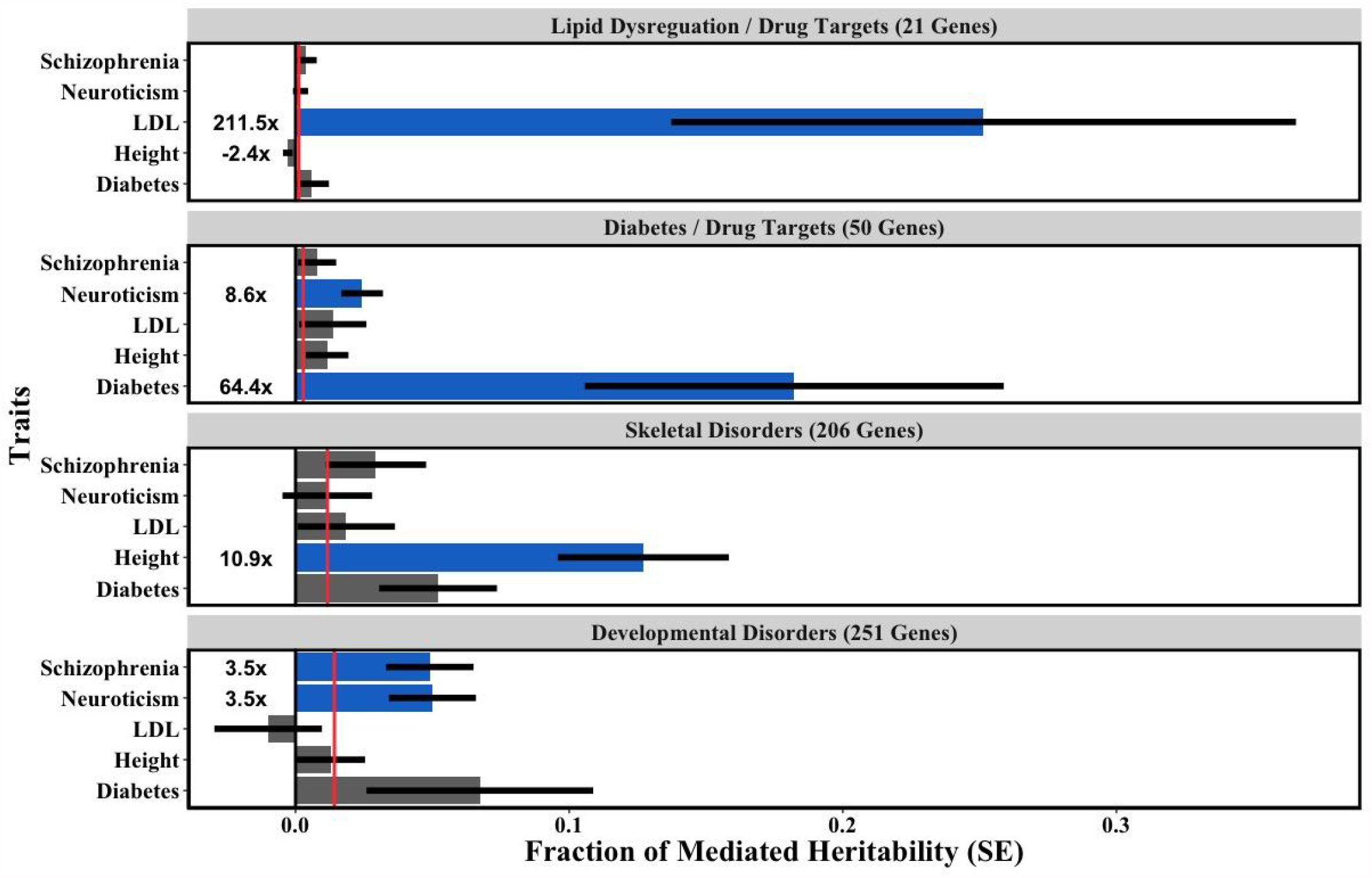
Mediated heritability enrichment of genes implicated in Mendelian disorders. Horizontal bars are the fraction of mediated heritability, calculated as the product of AMM enrichment and the fraction of total genes in the gene set (red line). For trait - gene set pairs with enrichments significantly greater than 1, the bars are colored blue and point estimate of enrichments listed. AMM enrichments estimates are also listed for enrichments significantly less than 1. Error bars represent standard errors of the fraction of mediated heritability. See Supplementary Table 1 for gene set references. For numerical results, see Supplementary Table 6.

Finally, we analyzed mediated heritability enrichment in 251 genes associated with developmental disorders (mostly neurodevelopmental). We find enrichments for cognitive traits in this gene set, including schizophrenia (3.5x, P = 0.01) and neuroticism (3.5x, P = 0.01). However, their enrichments are similar to constrained genes more broadly (respectively: 3.0x, 2.9x). The constrained gene set is approximately 10 times larger and thus mediates much more heritability than the neurodevelopmental gene set (47.4% vs. 4.9% in schizophrenia, for example). This modest enrichment could be related to the much higher polygenicity of brain-related traits compared with lipid and metabolic traits (see further discussion below).

### Mediated heritability enrichment of specifically expressed genes

SNPs near genes specifically expressed in trait-relevant tissues are enriched for heritability.^10^ However, it is unknown what fraction of heritability is actually mediated by these genes. We applied AMM to the top 10% of specifically expressed genes across GTEx tissues, as previously defined.^10^

AMM highlighted trait-relevant tissues (**Figure 5A**). Genes specifically expressed in cortex (vs. non-brain tissues) mediate 21.9% (SE: 3.5%, P = 2.7e-4) of schizophrenia heritability (2.2x enrichment), while genes specifically expressed in blood mediate 3.0% (SE: 3.5%, P = 0.02 for depletion). Genes specifically expressed in liver mediate 54.0% (SE: 18.4%, P = 8.3e-3) (5.4x enrichment) of LDL heritability. Genes specifically expressed in whole blood mediate 35.5% of Alzheimer’s disease heritability (SE: 12.3%, P = 0.02), while genes specifically expressed in cortex mediate 14.4% (SE: 7.5%, P = 0.28).

**Figure 5.**
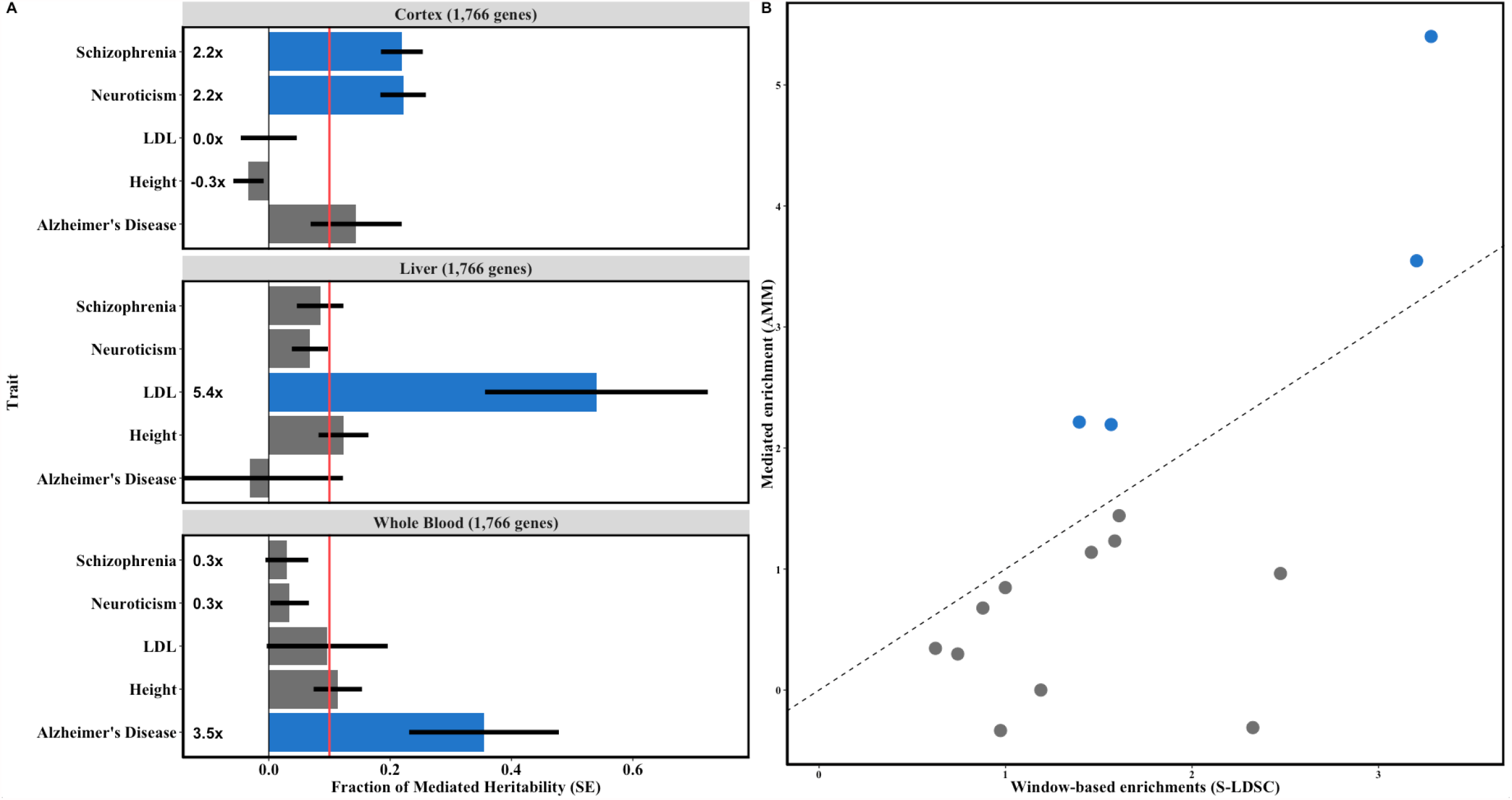
Mediated heritability enrichment of specifically expressed genes. **(A)** Horizontal bars are the fraction of mediated heritability, calculated as the product of AMM enrichment and the fraction of total genes in the gene set (red line). For trait - gene set pairs with enrichments significantly greater than 1, the bars are colored blue and point estimate of enrichments listed. For enrichments significantly less than 1, point estimates of enrichments are listed. Error bars represent standard errors of the fraction of mediated heritability. **(B)** Contrasting window-based enrichment (gene body +/- 100 kb) with mediated heritability enrichments (AMM) for the same 15 trait-gene set pairs as in Figure 5A (same color code). For numerical results, see Supplementary Tables 3-4, 7-8.

Using the same gene sets and traits, we estimated the heritability enrichment of SNPs within 100kb of these genes (“gene-window heritability”), defined as the fraction of heritability in the gene window divided by the fraction of SNPs, as described by Finucane et al. (**Figure 5B**).^10^ These estimates differ in two ways. First, gene-window enrichment estimates are attenuated relative to gene-mediated enrichments for significantly enriched trait-tissue pairs, as genes can be regulated by SNPs outside their window. Second, several trait pairs are depleted for mediated heritability but not for gene-window heritability. This difference results from the fact that gene-proximal SNPs are more likely to be functional (due to being coding, or conserved, etc), increasing their gene-window enrichment. This limitation affects point estimates of gene-window enrichment but not their p-values, which were defined more stringently (conditioning on a union-of-gene-windows annotation).^10^ AMM point estimates and p-values are unaffected by this phenomenon, as every SNP has exactly one closest gene, regardless of distance from it. In contrast to the point estimates, p-values for enrichment were similar between AMM and the window-based approach (Supplementary Figure 6).

### Contrasting mediated heritability enrichment of top expressed genes in cortex and liver

Among complex traits, the most polygenic are brain-related, and some of the least polygenic are liver-related.^32^ We investigated whether different gene expression patterns in the cortex compared with the liver from GTEx were related to these differences.^20^ First, we ranked genes by their fraction of (TPM normalized) expression in the focal tissue (see Online Methods) (**Figure 6A**). Expression in liver is skewed towards the most highly expressed genes: 50% of total expression in liver comes from the 131 most expressed genes, while in cortex the top 946 genes represent 50% of total expression. The most expressed gene in liver (*ALB*, albumin) represents 5.7% of total liver expression, compared to 0.7% for the most expressed gene in cortex.

**Figure 6.**
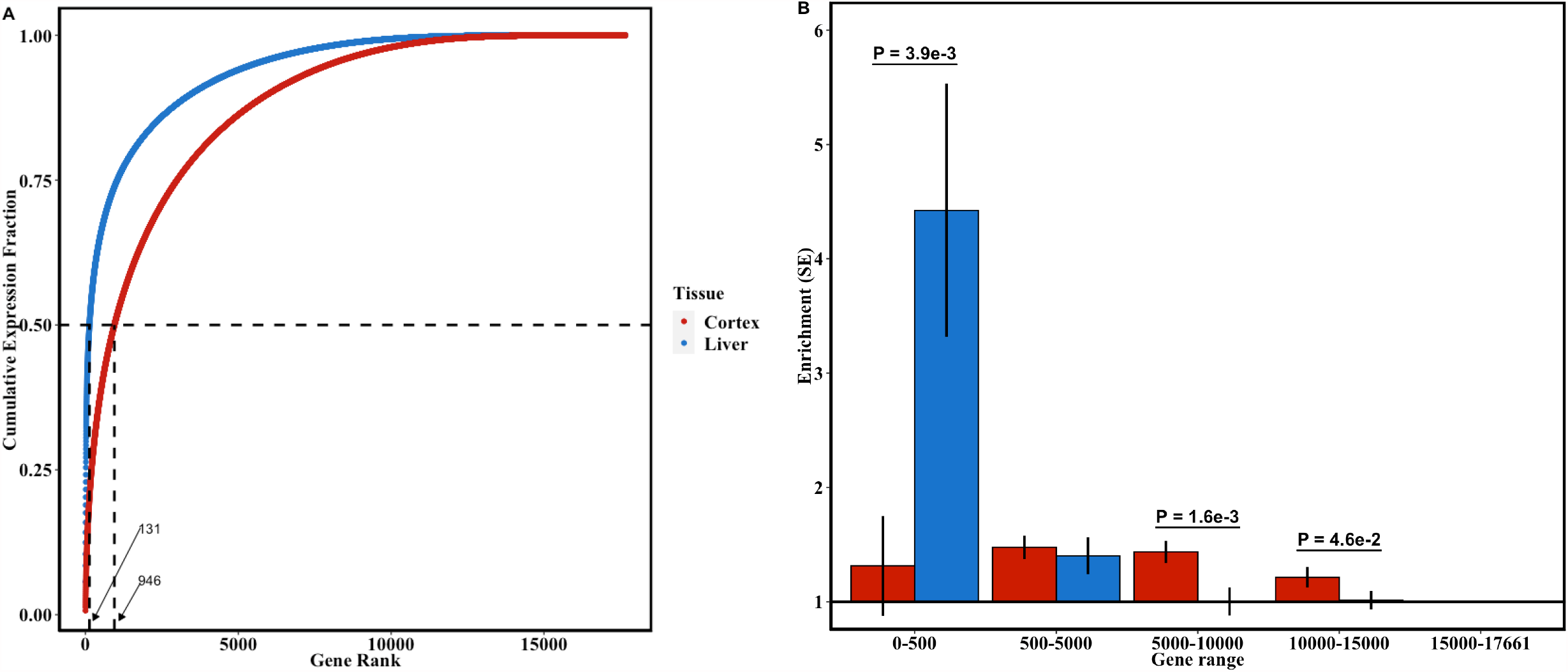
Contrasting expression and enrichment in cortex and liver. **(A)** We calculated the fraction of total expression for each gene as its median expression divided by the total across genes, and we plot the cumulative fraction of expression across genes from most to least expressed. **(B)** Across genes from most to least expressed, we calculated the cumulative mediated heritability enrichment of tissue-appropriate traits. Cumulative mediated enrichment for a bin was defined as its fraction of remaining heritability (subtracting more highly expressed genes) divided by its fraction of remaining genes; in the last bin, it is null by the definition. Error bars represent standard errors. For numerical results, see Supplementary Tables 9-10.

We estimated the cumulative mediated heritability enrichment for top expressed genes in each tissue, binning genes by expression from greatest to least expressed (**Figure 6B**). Cumulative mediated heritability enrichment was defined as the fraction of remaining heritability mediated by genes in the bin divided by the fraction of remaining genes, in a meta-analysis of tissue-enriched traits (see Online Methods and Supplementary Tables 3-4).

Like total expression, mediated heritability enrichment in liver is more highly concentrated among highly-expressed genes: the 500 most expressed genes were enriched in heritability (4.4x, P = 1.0e-3) and were significantly more enriched than the top 500 genes in cortex (P = 3.9e-3 for comparison). In contrast, the 5,000-10,000th and 10,000-15,000th most-expressed genes were enriched for heritability in cortex (P = 2.8e-6 and P = 8.4e-3, respectively) but unenriched in liver (P = 0.49 and P = 0.43, respectively; P < 0.05 for both comparisons). These estimates raise the question of whether differences in polygenicity among brain vs. liver-related traits are explained by differences in gene expression patterns.^33^

## Discussion

We have presented the Abstract Mediation Model for partitioning gene-mediated heritability and applied it to a set of 47 phenotypes and a range of biologically and biomedically important gene sets. Our results demonstrate that it is possible to estimate mediated heritability without observing underlying mediating variables, instead observing enriched proxies. This approach bypasses the need to measure gene expression levels in causal cell types and states.

Using AMM, we estimated that the closest gene mediates the most heritability (approximately 30%), and that a substantial fraction is mediated by genes outside the ten closest. This estimate is approximately concordant with the distances between significant eQTL - eGene pairs from GTEx (Supplementary Figure 2), as well as from previous estimates using eQTL and GWAS data.^17,27,28^ In contrast, a previous analysis of metabolite QTLs estimated that 69% of candidate causal genes were the closest gene to the lead variant.^34^ Our estimates of SNP-to-gene architecture are not directly comparable, as this previous analysis examined only genome-wide significant loci for metabolic traits, which have larger-than-average effect sizes. SNP-to-gene architecture may differ between loci with small and large effect sizes, for example if large-effect loci are disproportionately enriched in coding regions.

While we have utilized SNP-to-gene proximity to rank genes, other strategies could be considered. Hi-C contact,^35^ epigenetic co-expression^36^, and fine-mapped eQTL data,^37^ by themselves or together, could be used to rank genes for each SNP. The SNP-to-gene ranking should not be specifically informative for the gene set being tested (violating the “no-special-relationship” assumption; see Online Methods). An important goal should be to produce a maximally enriched SNP-to-gene mapping, such that the top ranked gene for each SNP explains the majority of disease heritability.

Our study has several limitations. First, AMM does not identify causal gene(s) for a given SNP, and more generally, putative disease gene sets must be prespecified. Second, AMM does not identify the cell type or context in which gene mediation occurs, potentially in contrast with eQTL-based approaches. Third, AMM often has limited power for individual trait - gene set pairs, requiring meta-analysis across traits. Fourth, AMM requires some assumptions about SNP-to-gene architecture (see Online Methods). Finally, stratified LD score regression has elevated false positive rates for small annotations, which may affect our analysis of small gene sets.^31^ Despite these limitations, this study presents a novel approach for estimating gene-mediated heritability and informs our understanding of the convergence of common and rare genetic variation. Finally, we have released software for implementing AMM at the command-line.

## Methods

### Abstract Mediation Model

The Abstract Mediation Model describes the expected effect-size variance of a SNP *x*_*i*_ on a trait in terms of (1) its effect-size variance on each nearby gene and (2) the effect-size variance of each gene on the trait. The SNP-to-gene effect-size variance depends on the proximity of the SNP to each gene (in particular, whether that gene is the closest, second-closest, etc. to each SNP). The gene-to-trait effect size depends on whether that gene is in a putatively enriched gene set.

Under the AMM, the effect-size variance of SNP *x*_*i*_ on a trait is:

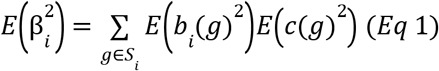

where

- β_i_ is the effect size of SNP *x*_*i*_ on the trait *Y*, in standardized units (SD of *Y* per SD of genotype)
- *b*_i_ (*g*) is the effect size of SNP *x*_*i*_ on gene *g*, in standardized units
- *c* (*g*) is the effect size of gene *g* on *Y*, in standardized units
- *S*_*i*_ is the set of nearby (cis) genes for SNP *x*

Let 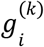 ∈ *S*_*i*_ be the *k*^*th*^ closest gene to SNP *x*_*i*_. The SNP-to-gene effect-size variance depends on the proximity rank of gene *g* with respect to SNP *x*_*i*_:

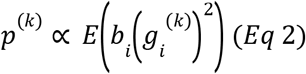

These *p*^*(k)*^ values are scaled to sum to one, and we interpret *p*^*(k)*^ as the proportion of *cis*-mediated heritability mediated by the *k*^*th*^ closest gene.

The gene-to-trait effect-size variance depends on whether the gene is in the gene set. We define:

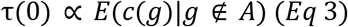

and

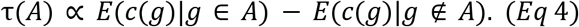

Substituting *p*^(k)^ τ(0) and τ(*A*) into Eq 1,

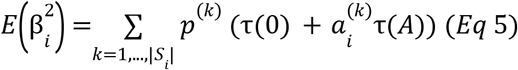

where 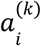 is a binary indicator of whether the *k*^*th*^ closest gene to SNP *x*_i_ is in A, and zero otherwise. This is the equation that is used for estimation (see below).

We define mediated heritability enrichment of a gene set as the average effect size variance of genes in A divided by the average effect size variance of all genes. If N(A)/N is the fraction of genes in A, then the mediated heritability enrichment of A, e(A), is given by:

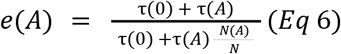

and the total proportion of heritability mediated by genes in A is defined as e(A)(N(A)/N). We note that this definition avoids assigning additional heritability to a gene set just because it has a larger number of nearby SNPs; we would view this source of enrichment as being spurious.

**Assumptions**. Estimation requires three primary assumptions:

1. **No-spurious-enrichment assumption**. We assume that the heritability enrichment of SNPs mapped to genes in A is actually mediated by those genes. This assumption is violated if genes in A lie in disease-relevant regions of the genome without being disease-relevant themselves. This assumption could be violated if A were chosen based on the GWAS itself: for example, suppose A is the set of genes which are the closest gene to a GWAS lead SNP. Even if none of these genes is actually the mediating gene, the set of SNPs whose closest gene is in A would be highly enriched for heritability, and we would wrongly infer large values for *e*(*A*) and *p*^(1)^. We do not apply AMM to gene sets that were constructed from GWAS data. This assumption also motivates us to condition on potential confounders in the baselineLD model.^25,26^ Some annotations may be enriched near genes in the gene set; for example, brain-expressed genes tend to be long, and their nearby SNPs may be enriched for the exonic and intronic annotations. It has previously been observed that conditioning on genomic annotations can explain certain gene-set enrichments.^38^
2. **No-special-relationship assumption**. We assume that cis-regulatory architecture, with respect to the cis-gene ranking, is the same for genes in *A* and genes not in *A*. More precisely, we assume that:

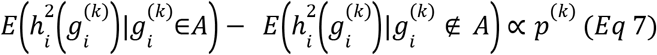

This assumption would be violated, for example, if A were the set of genes that are only regulated by promoter eQTLs and never by more distal enhancer eQTLs. It could also be violated if both the gene ranking and the gene set were cell-type-specific: for example, a ranking obtained from eQTL data in T cells may be highly informative for the set of T-cell-expressed genes, but not for B-cell-specific autoimmune disease genes. This source of model violations is not relevant to the proximity ranking (which is the same in all cell types).
3. **All heritability mediated assumption**. We assume that genes in *cis* mediate 100% of heritability. This assumption is plausible, as *trans*-regulatory effects are likely mediated by *cis*-regulatory effects^39^, but it could be violated if the number of *cis* genes were too small, or if some *trans*-regulatory effects are not mediated by any *cis* gene. Violations of this assumption would affect our estimates in two ways. First, estimates of *p*^(k)^are interpreted as proportions of *cis*-mediated heritability, rather than as proportions of all heritability. Second, our estimates of τ(0)would be upwardly biased, leading to downward bias in estimated heritability enrichments.

### Estimation

Let 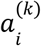 be the indicator variable for 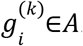. Suppose that genes inside and outside A have different effect-size variances, such that the heritability of SNP *x*_*i*_ mediated by its rank-*k* gene is:

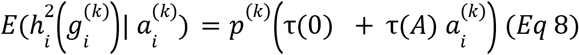

where τ(*A*) measures the heritability enrichment of genes in A and τ(0) measures the average heritability of all genes. This equation relies on assumption 2 (see above). Summing over genes,

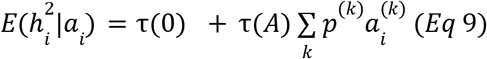

(We have used the definition that 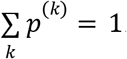) We can derive an LD score regression equation from Eq 9 (see Supplementary Note 1 for derivation). In brief, let 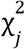 be the chi-squared statistic of SNP *x*_*j*_ calculated from a GWAS of size *N*, and let *l*_*j*_ (*A, k*) be the LD score for SNP *x*_*j*_ with the annotation 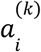 for all SNPs *x*_*i*_Then:

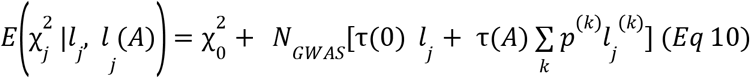

where *l*_*j*_ is the unstratified LD score and 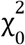 is the LD score regression intercept. This equation leads to the following estimation procedure:

1. Construct annotations 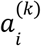 for each SNP and stratified LD scores for these annotations
2. Perform LD score regression on these LD scores jointly, possibly with additional covariates such as the baselineLD model (see below), obtaining regression coefficients 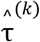 for each annotation
3. Estimate 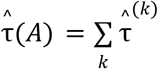
4. If 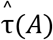 is significantly greater than zero, estimate 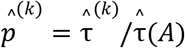

We use the output of Eq(10) to estimate the mediated heritability enrichment of the gene set A as noted above.

For all estimation, standard errors are derived from block jackknife with 200 partitions of adjacent SNPs, as previously described for LD score regression.^25^

Specific methods implemented in manuscript

### Estimation of *p*^*(k)*^ in bins

We binned gene proximity annotations to increase power. For instance, by binning the 3^rd^ through 5^th^ closest genes we obtained an average proportion of heritability mediated by each of the 3^rd^ through 5^th^ closest genes. We obtained binned *p*^*(k)*^ estimates through the following modified procedure:

1. Construct annotations 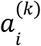 for each SNP and stratified LD scores for each of the *k* annotations. We defined gene location as the gene body midpoint.
2. For each bin range (i.e., 3^rd^ through 5^th^ closest genes), sum the LD scores for each annotation in the range, for example: 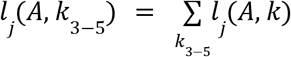
3. Perform LD score regression with these binned LD scores, plus additional covariates (such as baselineLD model)
4. Estimate 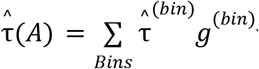 where g^(bin)^ is the number of genes in the bin.
5. Estimate binned 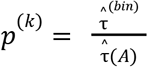

### Estimation of p^(k)^ with covariates and meta-analysis

We condition on baselineLD model annotations to eliminate potential sources of confounding, ensuring that geneset enrichments are not driven by genes that have larger numbers of nearby SNPs in one of these enriched annotations (for example, longer genes may have more nearby coding SNPs). The modified regression equation is:

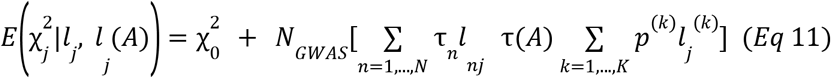

where τ_n_ is the regression coefficient for baselineLD model annotation *n* and *l*_*nj*_ is the LD score for SNP *j*.

When estimating *p*^*(k)*^ in constrained genes, we constructed a set of “baselineLD minus” annotations in order to avoid controlling for genic elements relevant to constrained genes; these excluded features include conservation, minor allele frequency, and ancient sequence annotation (Supplementary Table 11).

We estimated binned *p*^*(k)*^ using constrained genes and meta-analyzing across 47 traits. For each annotation bin (i.e. *k*^*th*^ closest gene is in the constrained set), we obtained binned estimates of τ^(k)^ in a joint regression as noted above and estimates of τ(A) by summing across binned annotations. To estimate binned p^(k)^ meta-analyzed across traits:

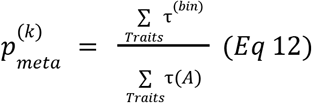

### AMM with known p^(k)^

AMM can simultaneously estimate p^(k)^ and mediated heritability enrichments for gene sets. However, we observed that power to estimate mediated heritability enrichments increases when pre-trained p^(k)^ are used (i.e. AMM only estimated mediated heritability enrichments and not p^(k)^ for the focal gene set) (Supplementary Figure 1). To use pre-trained *p*^*(k)*^, we use AMM to estimate a single heritability coefficient instead of a coefficient for each SNP annotation. To estimate this coefficient, we calculate a linear combination of LD scores for each of the original annotations, where the weights are the pre-trained *p*^*(k)*^ estimates. For instance, if analysis consisted of two annotations with pre-trained *p*^*(k)*^ values of [0.7, 0.3] with annotation LD scores *l*_1_ and *l*_2_, the new regression LD score would be *l*_pre-trained_ = 0.7*l*_1_ + 0.3*l*_2_. In the manuscript we used pre-trained *p*^*(k)*^ values from the set of constrained genes given its mediated heritability enrichment across a range of traits which facilitates meta-analysis.

We estimate τ(0) from the covariate annotations in the model (i.e. from the baselineLD model^26^). Specifically:

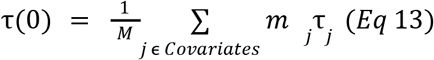

where *m*_*j*_ is the number of SNPs in the j^th^ covariate annotation, τ_j_ is the LD score regression coefficient for the j^th^ covariate annotation, and M is the number of SNPs in the model.

### GTEx gene sets

For analysis of mediated heritability of specifically expressed genes (Figure 5), we used previously defined gene sets based on bulk RNA expression across a range of tissues from GTEx.^10^ Briefly, in a given tissue, for each gene a t-statistic of specific expression as compared to other tissues was calculated. In a given tissue, specifically expressed genes are defined as the top 10% of genes with the greatest specific expression t-statistic. Note that for brain tissues (cortex) we used specific expression compared to non-brain tissues.

For analysis of mediated heritability enrichments of top expressed genes in cortex and liver (Figure 6), we downloaded from GTEx median gene-level TPM by tissue. The two focal tissues in this analysis were cortex (GTEx: ‘Brain - Cortex’) and liver (GTEx: ‘Liver’). We defined total expression in a focal tissue as the sum of per gene median expression across genes. We then calculated the fraction of total expression in a given tissue for a gene as the median expression of that gene in the tissue divided by the total expression in that tissue. We ranked genes by fraction of total expression in descending order and calculated cumulative expression as the fraction of total expression explained by that gene and all genes with greater total expression.

In Figure 6b, we estimated a modified version of enrichment, defined as the fraction of remaining heritability mediated in the bin divided by the fraction of remaining genes in the bin. In detail, for each gene bin, we used AMM to estimate a mediated heritability enrichment. For each bin, we then calculated the fraction of mediated heritability in that bin as the fraction of total genes in that bin multiplied by the mediated heritability enrichments. Starting at the first bin (Genes 1 - 1,000 by top expression), we then estimated the fraction of remaining heritability as the fraction of mediated heritability in the bin divided by fraction of mediated heritability explained by that bin and all bins of lower gene rank (i.e. lower mean expression).

### Window-based enrichments

Using sets of genes specifically expressed in tissues, we compared mediated heritability enrichments with window-based enrichments (Figure 5b). Estimation of window-based enrichments using stratified LD score regression followed the standard approach described previously.^10^ Briefly, we constructed SNP annotations +/- 100kb around the set of specifically expressed genes (see Online Methods: GTEx gene sets). We then estimated LD scores for this annotation and regressed this annotation and the Baseline model to control for potential confounding.^25^

### Consensus gene list

We defined a list of consensus genes to use as input for the kn-matrix. We constructed the consensus gene list as the intersection of (a) genes from the gnomAD browser (https://gnomad.broadinstitute.org/downloads) that were autosomal, had estimated pLI, and were protein-coding, (b) genes with a specific expression t-statistic from Finucane et al. 2018, and (c) genes with a median TPM measured from GTEx. The intersection of these three lists left 17,661 genes. Before analyzing any gene set, we first intersected the gene set with these 17,661 genes to ensure that all genes in the set were represented in our proximity matrix. We use GRCh37 in all analyses.

## Additional information

### Author contributions

D.J.W., S.G., E.B.R. and L.J.O designed the study. D.J.W. conducted the analyses. D.J.W. and L.J.O wrote the manuscript. D.J.W. wrote the AMM software.

## Acknowledgements

This work was supported by a grant from NLM (T15LM007092, D.J.W.) and by a grant from the Simons Foundation (704413, L.J.O. and E.B.R.). We are grateful to Jenna Ballard, Hilary Finucane, Sasha Gusev, Ajay Nadig, Alkes Price, Pouria Salehi, Katie Siewert, and Doug Yao for their input on this manuscript.

## Code Availability

Code to implement AMM at the command line and reference files for analysis are available at https://github.com/danjweiner/AMM21. All data used in these analyses is publically available.

