## Supplementary Figures and Note for "Partitioning gene-mediated disease heritability without eQTLs"

### **Supplementary Figures and Note for**
### ***Partitioning gene-mediated disease heritability without eQTLs***

**Supplementary Figures**

Supplementary Figure 1: Contrasting AMM enrichments and Z-scores for specifically expressed genes with and without pre-trained SNP-to-Gene architecture

Supplementary Figure 2: Distribution of significant eQTL-eGene proximity rankings, stratified by eGene constraint

Supplementary Figure 3: Enrichment estimates from the set of constrained genes

Supplementary Figure 4: Distribution of annotation correlations for constrained genes

Supplementary Figure 5: SNP-to-Gene architecture as a function of SNP-to-Gene distance

Supplementary Figure 6: Comparing mediated enrichment Z-scores from AMM with window-based coefficient Z-scores from S-LDSC

**Supplementary Note**

Supplementary Note 1: Derivation of AMM LD score estimator

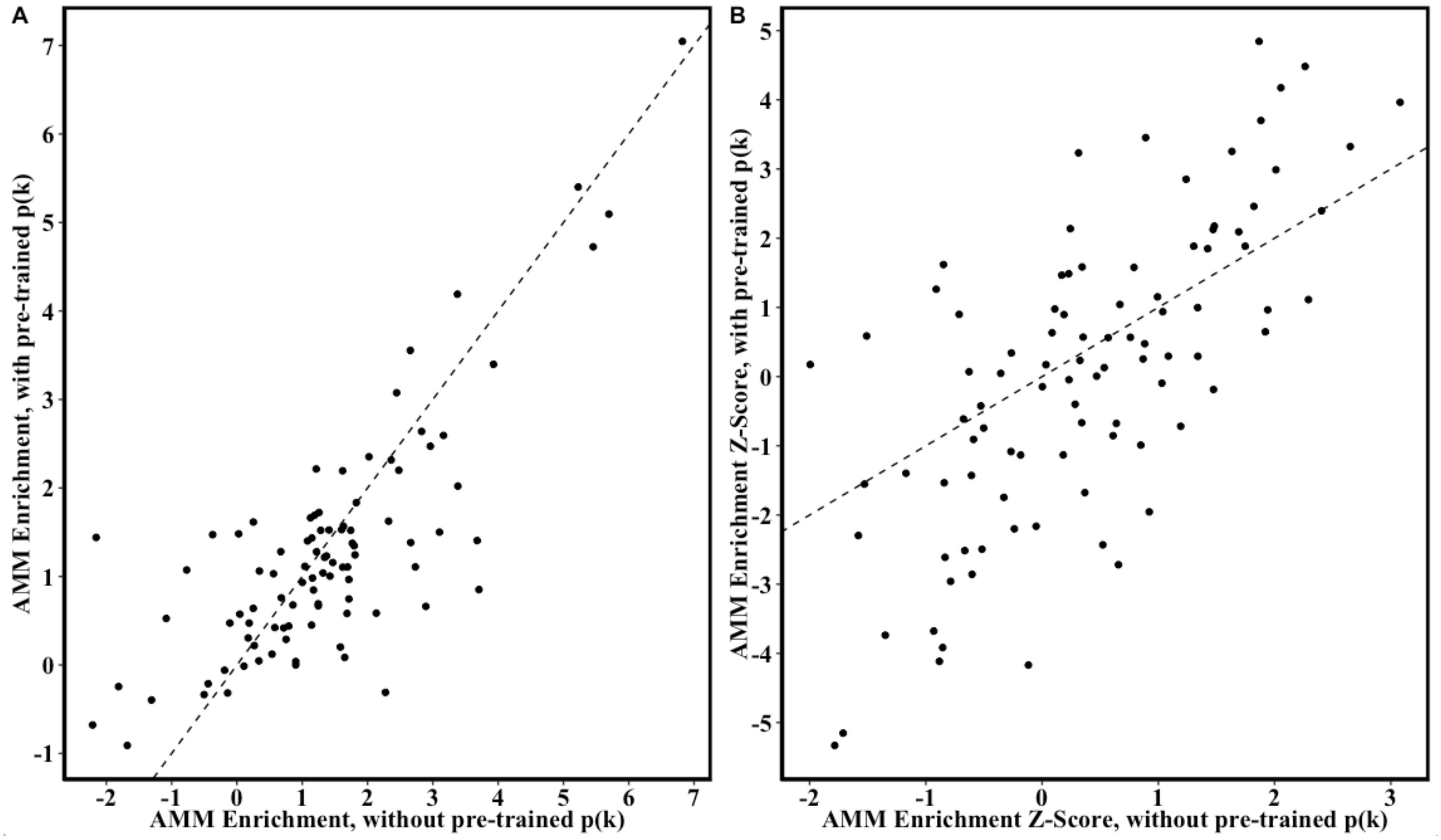


**Supplementary Figure 1: Contrasting AMM enrichments and Z-scores for specifically expressed genes with and without pre-trained SNP-to-Gene architecture**. (A) We calculated AMM enrichments for all 47 traits in the set of genes specifically expressed in liver and in cortex from GTEx (see Online Methods). For AMM enrichments without pre-trained $p^{(k)}$ (x-axis), we jointly estimated SNP-to-Gene architecture ($p^{(k)}$) and mediated heritability enrichments. For AMM enrichments with pre-trained $p^{(k)}$, we used SNP-to-Gene architecture estimates from the constrained gene set in the estimation of the mediated heritability enrichments (see Online Methods). (B) Same as (A), except contrasting AMM enrichment Z-scores.

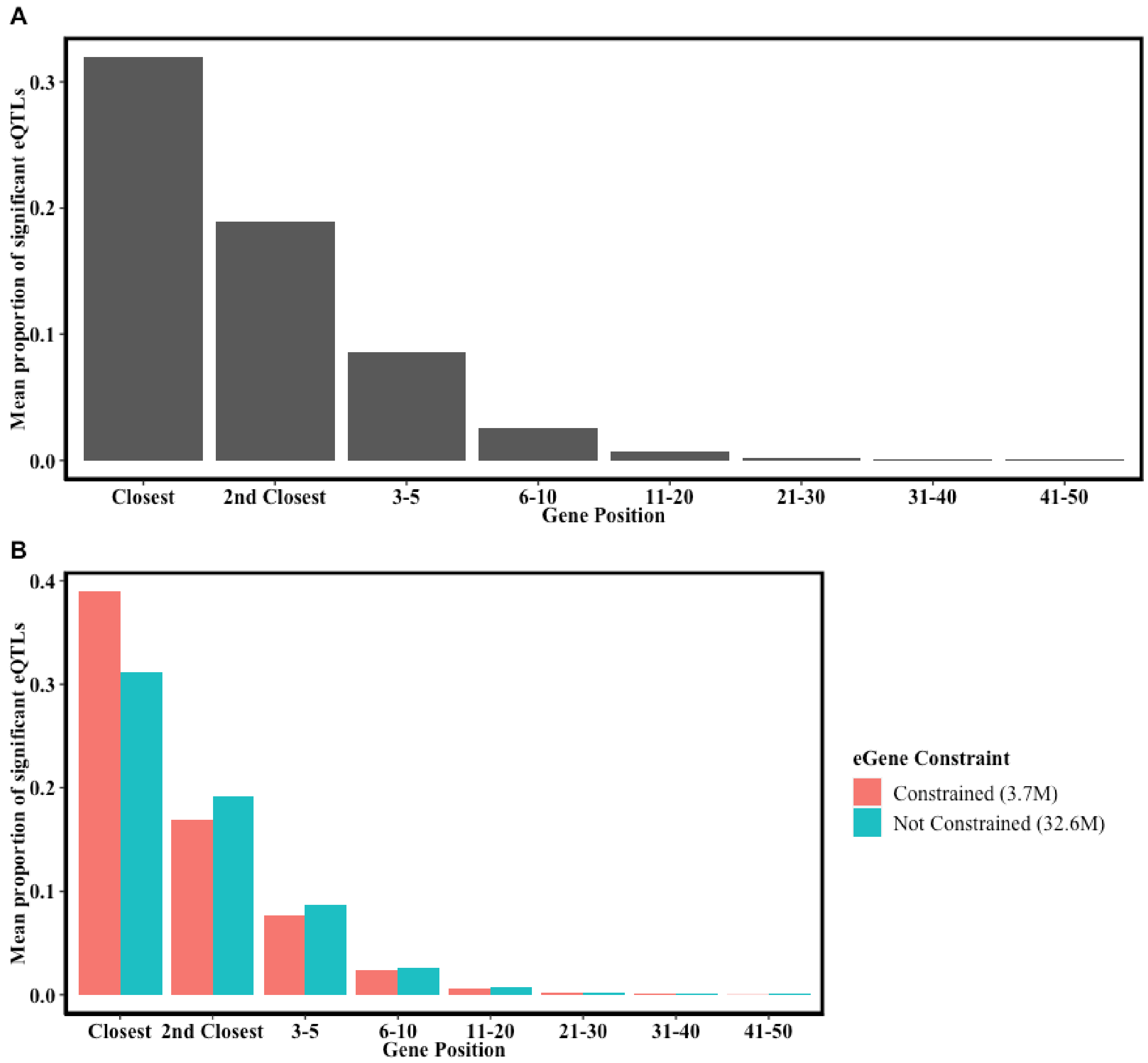


**Supplementary Figure 2: Distribution of significant eQTL-eGene proximity rankings, stratified by eGene constraint.** (A) We first identified all significant eQTL-eGene pairs from GTEx v8 across all 49 tissues. We then restricted to pairs whose eGene is more proximate than the 50th closest gene and has an estimated constraint metric pLI, leaving 36,287,362 eQTL-eGene pairs across all tissues. We then calculated the mean fraction of all eQTL-eGene pairs in each proximity bin. For example, 8.6% of all eQTL-eGene pairs have an eGene that is the 3rd - 5th closest gene to that eQTL. (B) As (A), stratified by whether the eGene is constrained (pLI $\geq$ 0.9).

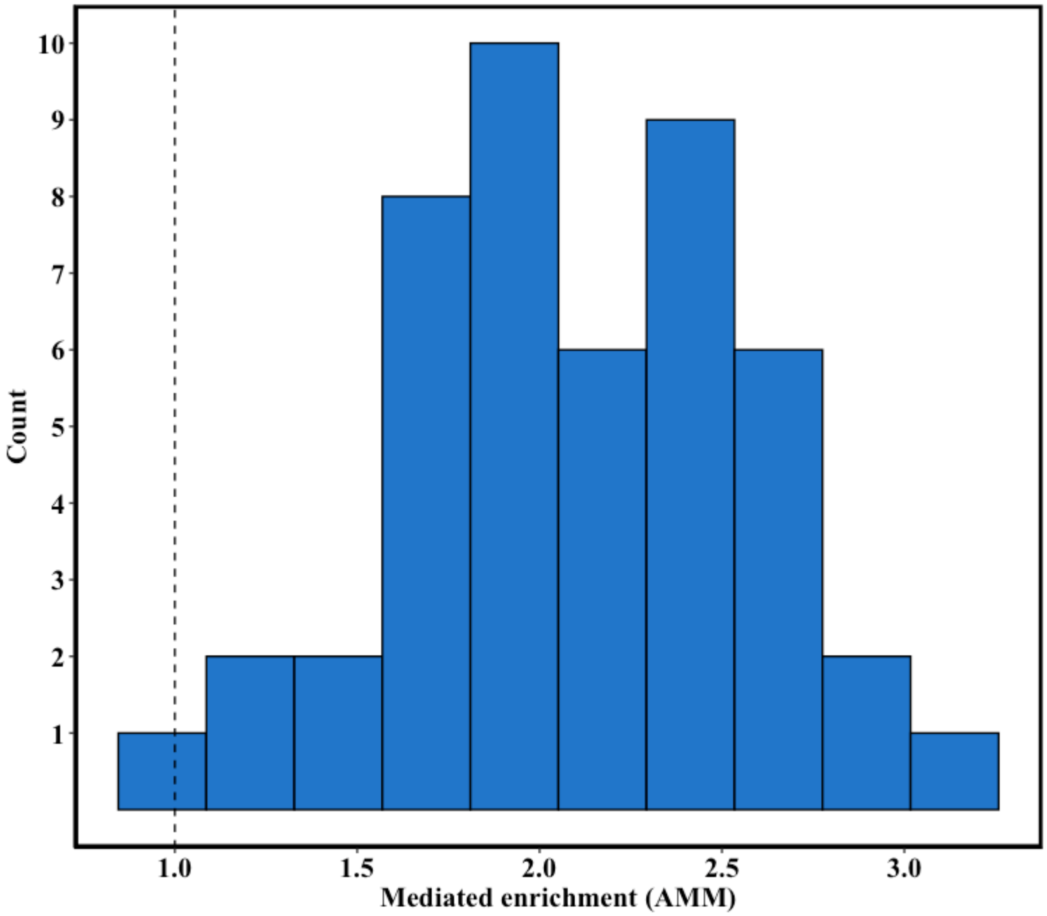


**Supplementary Figure 3: Enrichment estimates from the set of constrained genes.** Distribution of mediated heritability enrichment of 47 traits in the set of constrained genes. Median enrichment is approximately 2.1x; 46/47 traits with point estimate of enrichment greater than 1. Vertical dashed line at enrichment = 1 denotes threshold of positive enrichment.

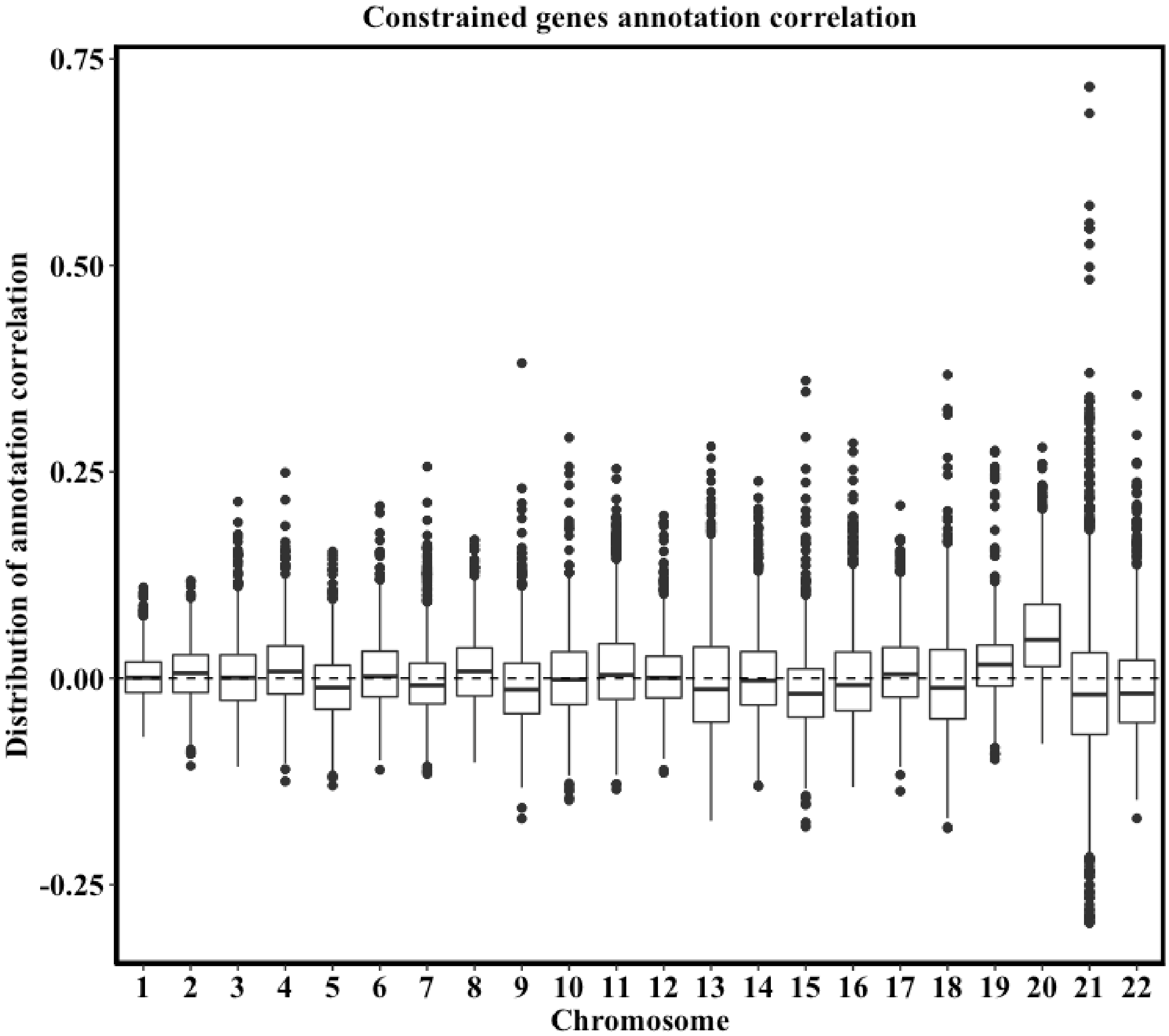


**Supplementary Figure 4: Distribution of annotation correlations for constrained genes**. We evaluated whether constrained genes cluster together through the correlations of their AMM annotations. If constrained genes clustered, then the gene proximity annotations would be correlated (i.e., correlation between the annotation for whether the closest and second closest gene to a SNP $x_i$ are constrained). Annotation correlation matrices were obtained from standard LD Score regression output. Upper and lower hinges represent 75th and 25th percentiles, respectively. Upper and lower whiskers represent 1.5*IQR from the respective hinge.

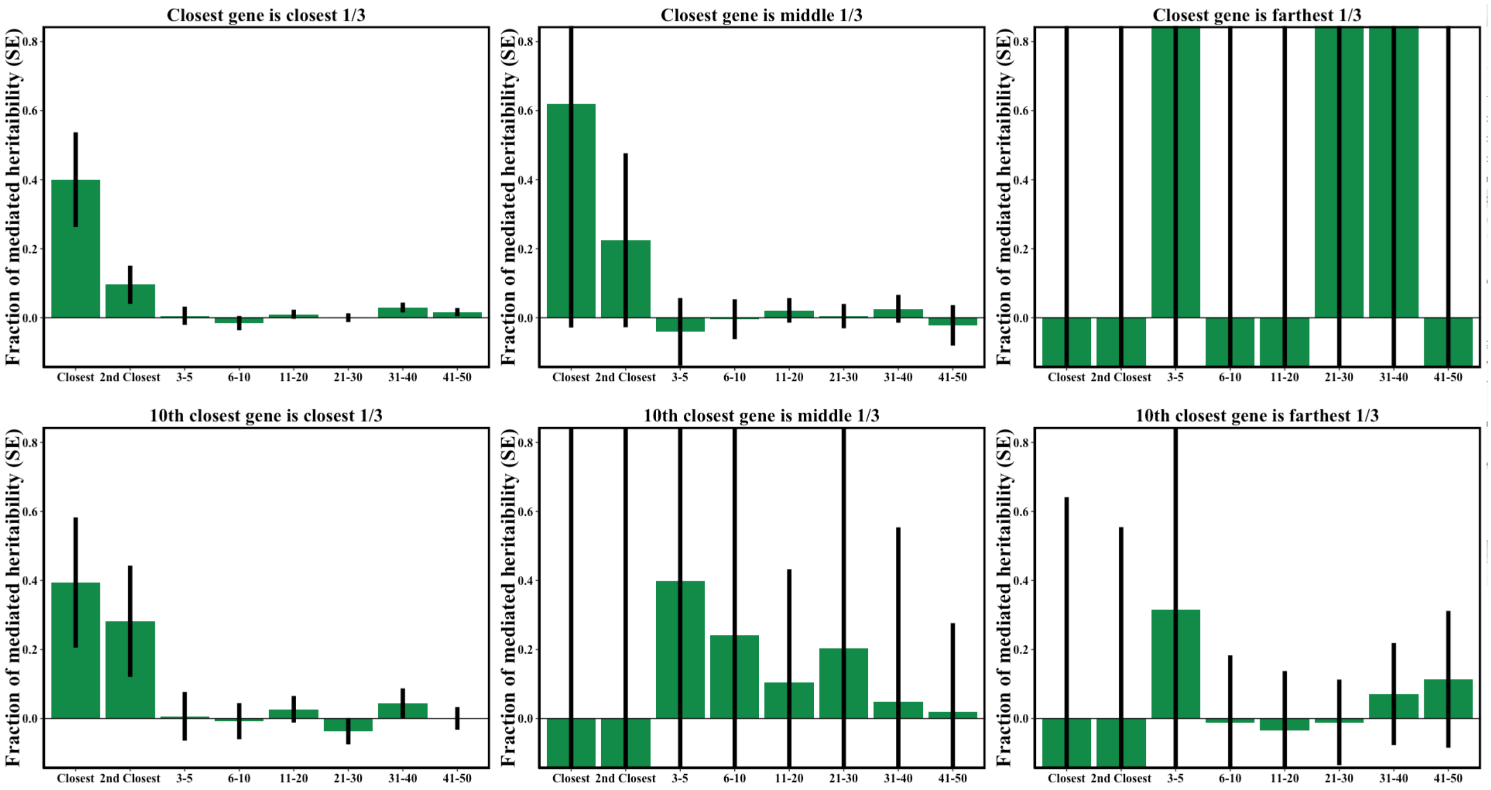


**Supplementary Figure 5: SNP-to-Gene architecture as a function of SNP-to-Gene distance.** We first calculated the distance between each SNP and each gene on the same chromosome (where gene location is defined as gene body midpoint). For each SNP, we then defined whether the distance to the $k^{th}$ closest gene to that SNP is in a particular distance quantile for that chromosome. For instance, starting on chromosome 1, we calculated the distribution of distances between SNPs and their closest genes, and binned SNPs according to their percentiles (i.e. closest third on the chromosome 1 distribution; within-chromosome binning was used to account for differences in gene density across chromosomes). This leads to a binary annotation for each SNP for whether its closest gene is in the closest third of SNP-Gene distance (and analogously for each middle and far third). We then multiplied this binary vector by the standard SNP-Gene rank matrix described in Online Methods, where the $k^{th}$ column vector is an annotation for whether the $k^{th}$ closest gene to a SNP is in a gene set A. The product is a matrix whose $k^{th}$ column vector is a binary annotation for whether the $k^{th}$ closest gene is in the gene set *and* is in the distance bin of interest (i.e. closest third). We then regressed the LD scores from this matrix, LD scores from the binary annotation across SNPs, and the baseline model on SNP effect size to generate estimates of $\tau(k)$.

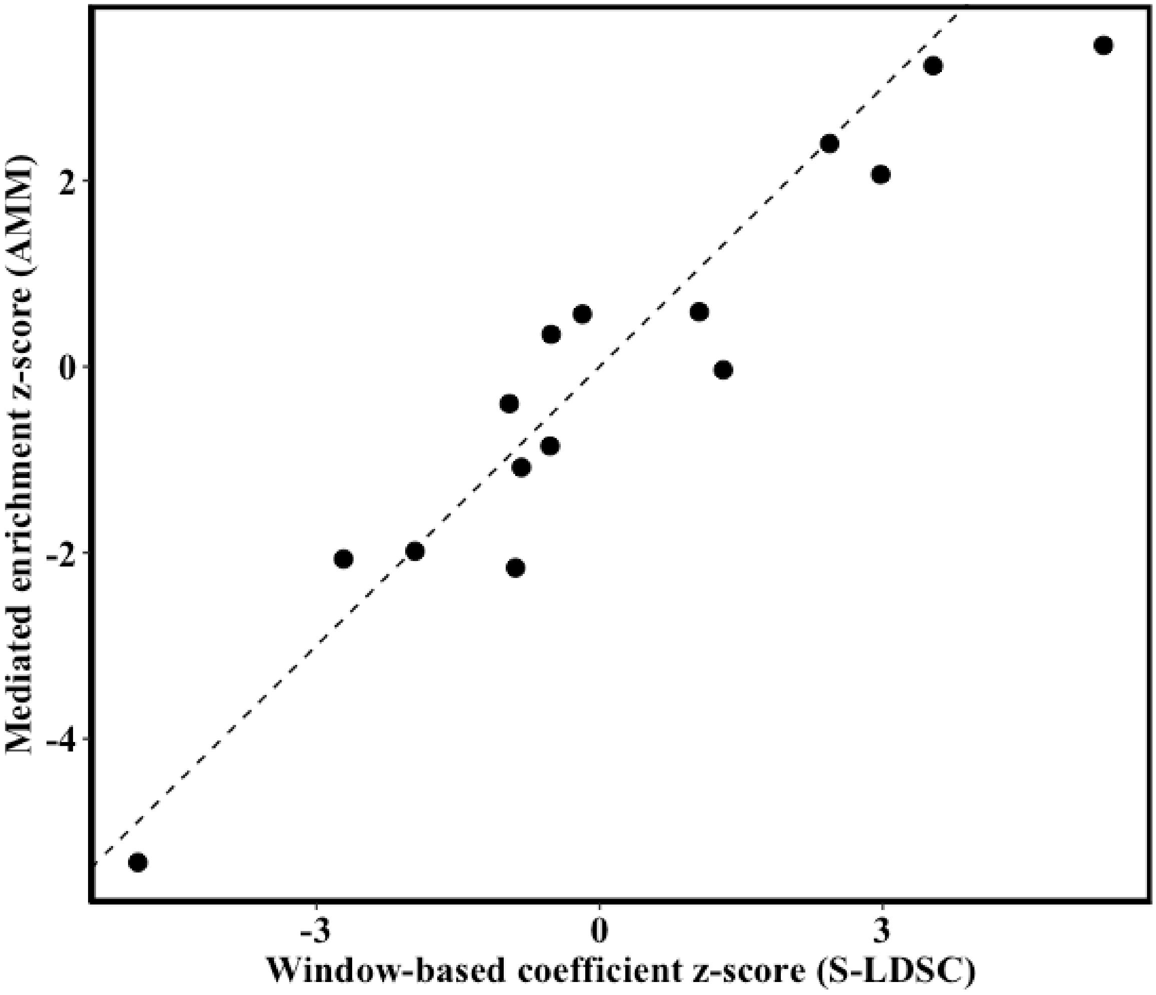


**Supplementary Figure 6: Comparing mediated enrichment Z-scores from AMM with window-based coefficient Z-scores from S-LDSC.** We compared the statistical power of mediated enrichments from AMM against window-based enrichments +/- 100kb. We used the same GTEx gene set - trait pairs from Figure 5A-B. The AMM enrichment z-scores are defined as (*e(A)* - 1)/SE(*e(A)*). The S-LDSC z-score is from the regression coefficient of annotation of specifically expressed genes.

**Supplementary Note 1**: Derivation of AMM LD score estimator

Our goal is to develop the following expression for per SNP heritability into an LD score regression equation:

$$E(h_i^2|a_i) = \tau(0) + \tau(A)\sum_k p^{(k)} a_i^{(k)} \quad (Eq\ S1)$$

Let $E[h_i^2|a_i] = E[\beta_i^2]$, $\tau(k) = \tau(A)p^{(k)}$, define $a^0_i = 1$ for all SNPs $x_i$, and let |S| be the number of genes in *cis*. Thus:

$$E[\beta_i^2] = \tau(0)a_i^0 + \sum_{k=1}^{|S|} \tau(k)a_i^{(k)} \text{(Eq S2)}$$

$$E[\beta_i^2] = \sum_{k=0}^{|S_i|} \tau(k)a_i^{(k)} \text{(Eq S3)}$$

The expected marginal effect size of SNP $x_j$, $\alpha_j$, is related to its squared correlation $r_{ij}^2$ with the causal effect sizes, $\beta_i$ of other SNPs $x_i$:

$$E[\alpha_j^2] = \sum_i r_{ij}^2 E[\beta_i^2] \text{(Eq S4)}$$

Substituting Eq S4 into Eq S3 yields:

$$E[\alpha_j^2] = \sum_i r_{ij}^2 \sum_{k=0}^{|S|} \tau(k)a_i^{(k)} \text{(Eq S5)}$$

By summation rearrangement:

$$E[\alpha_j^2] = \sum_{k=0}^{|S|} \tau(k) \sum_i r_{ij}^2 a_i^{(k)} \text{(Eq S6)}$$

We define the stratified LD score of SNP $x_j$ to annotation $k$, $l_j^{(k)}$, as the sum of the squared correlation between SNP $x_j$ and all SNPs $x_i$ in annotation k:

$$l_j^{(k)} = \sum_i r_{ij}^2 a_i^{(k)} \quad (Eq\ S7)$$

Substituting Eq S7 into Eq S6 yields:

$$E[\alpha_j^2] = \sum_{k=0}^{|S|} \tau(k) l_j^{(k)} \text{(Eq S8)}$$

Given marginal $\chi^2$ statistics from GWAS, we can express Eq S8 in LD score regression form:

$$E[\chi_j^2] = 1 + N_{GWAS} \sum_{k=0}^{|S|} \tau(k) l_j^{(k)} \text{(Eq S9)}$$

We define the LD score of SNP $x_j$ to all SNPs, $l_j$ as the unstratified LD score, and expand the summation in Eq S10 to yield:

$$E[\chi_j^2] = 1 + N_{GWAS}[\tau(0) l_j + \sum_{k=1}^{|S|} \tau(k) l_j^{(k)}] \text{(Eq S10)}$$

Using the definition $\tau(k) = \tau(A)p^{(k)}$ again yields our desired LD score regression equation:

$$E[\chi_j^2] = 1 + N_{GWAS}[\tau(0) l_j + \tau(A) \sum_{k=1}^{|S|} p(k) l_j^{(k)}] \text{(Eq S11)}$$
